# Association between ocular biometrical parameters and diabetic retinopathy in Chinese adults with type 2 diabetes mellitus

**DOI:** 10.1101/2020.02.06.937144

**Authors:** Lanhua Wang, Sen Liu, Wei Wang, Miao He, Zhiyin Mo, Xia Gong, Kun Xiong, Yuting Li, Wenyong Huang

**Affiliations:** Zhongshan Ophthalmic Center, State Key Laboratory of Ophthalmology, Sun Yat-Sen University, Guangzhou, People’s Republic of China; School of Medicine, Sun Yat-sen University Guangzhou, China; Department of Ophthalmology, Guangdong General Hospital, Guangdong Academy of Medical Sciences, Guangzhou, People’s Republic of China

**Keywords:** diabetic retinopathy, myopia, axial length, AL/CR ratio, ocular biometry

## Abstract

**Purpose:** To investigate the association between ocular biometrical parameters and diabetic retinopathy (DR) in ocular treatment naive patients with diabetes.

**Methods:** This cross-sectional study recruited type 2 diabetes mellitus patients with no history of ocular treatment in Guangzhou, China. The ocular biometrical parameters were obtained by Lenstar, including corneal diameter, central corneal thickness (CCT), corneal curvature (CC), anterior chamber depth (ACD), lens thickness (LT), and axial length (AL). The lens power and axial length-to-cornea radius ratio (AL/CR ratio) were calculated. Spherical equivalent (SE) was determined by auto-refraction after pupil dilation. Multivariate logistic regression analyses were performed to explore the associations of ocular biometry with any DR and vision threatening DR (VTDR).

**Results:** A total of 1838 patients were included in the final analysis, involving 145 5(79.2%) patients without DR and 383(20.8%) patients with DR. After adjusting confounding factors, any DR was independently associated with AL (OR = 0.84 per 1 mm increase, 95%CI: 0.74, 0.94), lens power (OR = 0.9951 per 1 D increase, 95%CI: 0.9904, 0.9998), and AL/CR ratio (OR = 0.26 per 1 increase, 95%CI: 0.10-0.70). Similarly, the presence of VTDR was independently related to AL (OR = 0.67 per 1 mm increase, 95%CI: 0.54-0.85), lens power (OR = 0.99 per 1 D increase, 95%CI: 0.98, 0.997), and AL/CR ratio (OR = 0.04 per 1 increase, 95%CI: 0.01, 0.25). The CC, corneal diameter, and refractive status were not significantly correlated with presence of DR or VTDR.

**Conclusion:** Longer AL, deeper ACD, higher lens power, and higher AL/CR ratio may be protective factors against DR. Considering the high prevalence of myopia in the Chinese juvenile population, it is worth paying attention to how the incidence of DR in this generation may change over time.

## Introduction

Diabetic retinopathy (DR) is a common cause of visual impairment in the working-age population.^1^ However, the pathogenesis of DR still remains unclear. Systemic risk factors for DR (e.g. course of diabetes, blood glucose and blood pressure) have broadly drawn the attention of investigators; however, few studies have focused on ocular risk factors.^2^ Prevalence of myopia increased significantly in the last decades globally, especially become epidemic in some Asian regions.^3^ It was found however in clinical practice that diabetic patients with myopia are less likely to suffer from severe DR.^4^ A small number of clinical studies and epidemiological studies have suggested that myopia could be a protective factor against DR; however, this conclusion remains controversial.^4–7^

Whether the ocular structure or the refracting media, or both, contribute to the protective relationship is still under debate. Several studies have evaluated the association of axial length (AL), myopia and refracting media in cases of DR.^6–10^ However, the investigators have not yet reached a consensus on the implications of these results. Similarly, the association of the anterior chamber depth (ACD), lens, cornea curvature (CC) and AL/CR ratio with the risk of DR was elusive.^9, 11^ Furthermore, the majority of previous studies were conducted no further subdivision to exclude the influence of potential confounding factors including age, sex, glycaemic level, and history of cataract surgery, which may bias the results.^12^ Thus, the which component of the ocular biometrical parameter contribute to the association between myopia and DR remains unclear.

The influence of myopia and ocular biometry parameters on the occurrence and progression of DR needs further clarification; therefore, the objective of this study was to investigate the association between ocular biometrical parameters and DR assessed in a large sample size of patients with type 2 diabetes mellitus (T2DM).

## Methods

### Subjects

This cross-sectional study was performed at Zhongshan Ophthalmic Centre (ZOC), Sun Yat-sen University, China. The protocol of the study was approved by the Institute Ethics Committee of ZOC and the study was performed according to the guidelines of the Helsinki Declaration. All subjects gave their informed written consent prior to enrolling in the study. Individuals aged 30 to 85 who were diagnosed with T2DM without ocular treatment history were recruited from the local government diabetes registry system. Exclusion criteria were as follows: (1) history of serious systemic diseases except for diabetes, such as serious cardiovascular and cerebrovascular diseases, malignant tumour, renal diseases; (2) history of systemic surgery such as coronary artery bypass graft, thrombolytic therapy and renal transplant; (3) presence of cognitive impairment, mental disorders, or inability to complete the questionnaire and examination; (4) history of ocular surgical interventions, such as retinal laser, intraocular injection, glaucoma surgery, cataract surgery, or laser myopia surgery; and (5) abnormal refractive media (severe cataract, corneal ulcer, pterygium, or corneal turbidity), poor fixation and other conditions resulting in poor quality of the fundus images.

### Procedures and definitions

A detailed questionnaire was used to collect the subjects’ demographic data, lifestyle risk factors, medical history and medication history. Outpatient medical records were reviewed to confirm details of medical history. All subjects underwent complete ocular examinations, including visual acuity, intraocular pressure (IOP), slit lamp examination, fundus examination, auto refraction, ocular biometric measurement, fundus photography, OCT and OCTA. The right eye was examined first in all eye examinations. Best corrected vision acuity (BCVA) were measured using ETDRS LogMAR E charts (Precision Vision, Villa Park, Illinois, USA) at a distance of 4 m. The IOP was measured with a noncontact tonometer (CT-1 Computerized Tonometer, Topcon Ltd., Topcon). The anterior and posterior segments were examined with a slit lamp bio-microscope (BQ-900, Haag-Streit, Switzerland) and a 90 D indirect ophthalmoscope.

Refraction errors were measured with an auto refractometer (KR-8800; Topcon, Japan), and spherical equivalent (SE) was calculated by adding half of the cylindrical power to the spherical power. Emmetropia was defined as SE between −0.5 and 0.5 D. Myopia, mild myopia, moderate myopia, and high myopia were defined as SE <−0.5 D, −0.5 to −3 D, −3 to −6 D, and < −6 D, respectively. Hyperopia was defined as SE >0.5 D.^13^

Ocular biometric parameters were measured using Lenstar LS900 (Haag-Streit AG, Koeniz, Switzerland), including central corneal thickness (CCT), corneal diameter, corneal curvature (CC), anterior chamber depth (ACD), lens thickness (LT), axial length (AL). The axial length-to-corneal radius ratio (AL/CR ratio) was defined as the AL divided by the mean radius of the corneal curvature. The lens power was calculated according to the modified Bennette-Rabbetts formula.^14^

Pupil dilation was performed with instillation of 0.5% tropicamide plus 0.5% phenylephrine eye drops. Once the pupils were fully dilated, standardised 7-field colour retinal photographs were taken, according to criteria from the Early Treatment Diabetic Retinopathy Study (ETDRS), using a digital fundus camera (Canon CR-2, Tokyo, Japan). Diabetic retinopathy (DR) was graded according to the American Academy of Ophthalmology (AAO) International Clinical Diabetic Retinopathy Disease Severity Scale by 2 ophthalmologists.^15^ Any DR was defined as presence of non-proliferative DR (NPDR), proliferative DR (PDR), diabetic macular oedema (DME), or any combination; and vision threatened DR (VTDR) was defined as presence of PDR and / or DME.^16^

Systolic and diastolic blood pressure were measured using a blood pressure monitor (HBP-9020; OMRON, Osaka, Japan). Weight and height were measured using an automatic weight and height scale (HNH-318; OMRON), with subjects standing on the scale with light clothes and no shoes on. All examinations were performed in compliance with the standardisation of procedures by an experienced nurse. Body mass index (BMI) was calculated as the weight (kg) divided by the square of height (m). Fasting (8 hours) venous blood samples were collected and sent for analysis of serum creatinine (Scr), glycosylated haemoglobin (HbA1c), total cholesterol (TC), high-density lipoprotein cholesterol (HDL-c), low-density lipoprotein cholesterol (LDL-c), triglyceride (TG), serum uric acid, and urine microalbuminuria (MAU) in accordance with standardised procedures from a certified laboratory in China.

### Statistical analyses

Only the data of worse eyes were incorporated into the analyses. The Kolmogorov-Smirnov test was conducted to verify normal distribution. When normality was confirmed, a t-test was carried out to evaluate the differences in demographic, systemic and ocular parameters between patients with and without DR. Next, Pearson’s correlation analyses were conducted to assess the associations between ocular components and HbA1c. Univariate and multivariate logistic regression analyses were used to explore the correlations of ocular biometry and diabetic retinopathy in patients with any DR or with (VTDR). Potential cofounding factors were first adjusted for age and sex, and then adjusted for age, sex, course of T2DM, HbA1c, TC, serum Scr, serum uric acid, BMI and mean blood pressure. We next investigated the dose-response relationship between AL as a continuous variable and presence of DR or VTDR. We used restricted cubic splines with 5 knots located at 5%, 27.5%, 50%, 72.55 and 95% of the distribution of AL. All analyses were performed using Stata Version 14.0 (Stata Corporation, College Station, TX, USA). P value of < 0.05 was considered statistically significant.

## Results

### Demographic and clinical characteristics

A total of 1838 individuals were included in the final statistical analyses, involving 145 5(79.2%) patients without DR and 383(20.8%) patients with DR. Table 1 shows the demographic and clinical characteristics of the subjects. The mean age of the subjects was 64.5 ± 7.9 years, 42.7% of them were male, and the duration of diabetes was 9.0 ± 7.9 years. Of the 1455 patients without DR, 591(40.6%) were male, the average age was 64.6±8.0 years, and the average course of diabetes was 8.3±6.7 years. Of the 383 patients with DR, 193(50.4%) were male, the average age was 64.1±7.9 years and the duration of diabetes was 11.7±7.5 years. Demographically, individuals with DR had a longer course of diabetes (P < 0.001). In terms of systemic and ocular parameters, subjects with DR also had higher levels of HbA1c, higher mean blood pressure, higher serum creatinine level, shorter AL, smaller lens power, and smaller AL/CR ratio (all, P < 0.05). There were no differences in age, BMI, total cholesterol, TG, serum uric acid, CCT, CC, corneal diameter, CCT, ACD and LT (all, P > 0.05) between individuals with and without DR.

**Table 1.** Demographic and clinical features of the included patients with type 2 diabetes mellitus.

|  | **Overall** | **Non-DR** | **Any-DR** | **P-value** |
| --- | --- | --- | --- | --- |
| No. of subjects | 1838 | 1455(79.2%) | 383(20.8%) | - |
| Gender |  |  |  | **0.001** |
| Male | 784(57.3%) | 591(40.6%) | 193(50.4%) |  |
| Female | 1054(42.7%) | 864(59.4%) | 190(49.6%) |  |
| Mean age, year | 64.5±7.9 | 64.6±8.0 | 64.1±7.9 | 0.239 |
| Duration of diabetes, year | 9.0±7.0 | 8.3±6.7 | 11.7±7.5 | **<0.001** |
| HbA1c, % | 7.0±1.4 | 6.8±1.3 | 7.7±1.7 | **<0.001** |
| Body mass index, kg/m<sup>2</sup> | 24.7±3.3 | 24.7±3.3 | 24.5±3.2 | 0.306 |
| Mean BP, mmHg | 182.1±23.6 | 180.7±23.2 | 187.3±24.2 | **<0.001** |
| Total cholesterol, mmol/L | 4.8±1.0 | 4.8±1.0 | 4.8±1.1 | 0.548 |
| Triglycerides, mmol/L | 2.3±1.6 | 2.3±1.6 | 2.3±1.7 | 0.630 |
| Serum creatinine, µmol/L | 72.9±22.7 | 71.4±20.1 | 78.6±30.1 | **<0.001** |
| Serum uric acid, µmol/L | 367.4±97.6 | 367.6±96.4 | 366.7±102.2 | 0.866 |
| Spherical equivalent, D | 0.1±4.5 | -0.04±2.7 | 0.1±2.4 | 0.247 |
| CCT, µm | 546.8±31.7 | 546.4±31.7 | 548.1±31.7 | 0.345 |
| Corneal diameter, mm | 11.6±0.4 | 11.6±0.4 | 11.6±0.4 | 0.269 |
| Corneal curvature, D | 44.2±1.5 | 44.2±1.5 | 44.2±1.5 | 0.733 |
| Axial length, mm | 23.6±1.2 | 23.6±1.2 | 23.4±1.2 | **0.006** |
| ACD, mm | 2.4±0.4 | 2.4±0.4 | 2.4±0.4 | 0.199 |
| Len thickness, mm | 4.7±0.3 | 4.7±0.3 | 4.7±0.4 | 0.112 |
| Lens power, D | -144.3±30.9 | -143.4±31.8 | -147.9±26.7 | **0.014** |
| AL/CR ratio | 3.08±0.14 | 3.09±0.15 | 3.07±0.13 | **0.005** |

Data were presented as mean ± standard deviation (SD).
DR=diabetic retinopathy; BP=blood pressure; CCT=central corneal thickness; ACD=anterior
chamber depth; AL/CR ratio=axial length-to-corneal radius ratio.
Bold indicates statistical significance.

### Ocular biometrical components and HbA1c

Table 2 shows the correlation between different ocular biometrical components and HbA1c levels. Both CCT and lens power were correlated with HbA1c (r = 0.0839, P = 0.0014 for CCT, r = −0.0669, P = 0.0051 for lens power), respectively. Other ocular parameters showed no correlation with HbA1c, including SE, corneal diameter, CC, AL, ACD, LT, and AL/CR ratio (all, P > 0.05). AL was correlated with all other ocular biometrical parameters (all, P < 0.05).

**Table 2.** Correlation coefficients between ocular components and HbA1c levels.

| SE |  |  |  |  |  |  |  |  |  |
| --- | --- | --- | --- | --- | --- | --- | --- | --- | --- |
| r=-0.0359<br>P=0.1305 | CCT |  |  |  |  |  |  |  |  |
| r=-0.0422<br>P=0.0758 | r=-0.0011<br>P=0.9619 | Corneal<br>diameter |  |  |  |  |  |  |  |
| r=-0.0276<br>P=0.2451 | r=-0.1608*<br>**P<0.001** | r=-0.4953*<br>**P<0.001** | Corneal<br>curvature |  |  |  |  |  |  |
| r=-0.4493*<br>**P<0.001** | r=0.0889*<br>**P<0.0012** | r=0.3880*<br>**P<0.001** | r=-0.4577*<br>**P<0.001** | AL |  |  |  |  |  |
| r=-0.1722*<br>**P<0.001** | r=-0.0606*<br>**P=0.0098** | r=0.3090*<br>**P<0.001** | r=-0.0076<br>P=0.7477 | r=0.3882*<br>**P<0.001** | ACD |  |  |  |  |
| r=0.1266*<br>**P<0.001** | r=-0.0017<br>P=0.9417 | r=-0.0396<br>P=0.0949 | r=-0.0346<br>P=0.1443 | r=-0.1752*<br>**P<0.001** | r=-0.5890*<br>**P<0.001** | Len<br>thickness |  |  |  |
| r=0.5889*<br>**P<0.001** | r=-0.1205*<br>**P<0.001** | r=0.1244*<br>**P<0.001** | r=0.1183*<br>**P<0.001** | r=0.0151<br>P=0.5264 | r=0.4523*<br>**P<0.001** | r=-0.1624*<br>**P<0.001** | Lens power |  |  |
| r=-0.5101*<br>**P<0.001** | r=-0.0207<br>P=0.379 | r=0.0648*<br>**P=0.0058** | r=0.2204*<br>**P<0.001** | r=0.7656*<br>**P<0.001** | r=0.4199*<br>**P<0.001** | r=-0.2167*<br>**P<0.001** | r=0.1048*<br>**P<0.001** | AL/CR<br>ratio |  |
| r=-0.021<br>P=0.3777 | r=0.0839*<br>**P<0.0014** | r=-0.0062<br>P=0.7925 | r=-0.0022<br>P=0.9254 | r=-0.0325<br>P=0.1676 | r=0.0028<br>P=0.9053 | r=-0.0249<br>P=0.2954 | r=-0.0669*<br>**P=0.0051** | r=-0.0366<br>P=0.1209 | HbA1c |

\*Bold indicates statistical significance.
SE=spherical equivalent; CCT=central corneal thickness; AL=axial length; ACD=anterior chamber depth; AL/CR ratio=axial length to corneal curvature ratio.

### Ocular biometry parameters and DR presence

Table 3 presents the association between ocular biometry and DR after adjusting for age and sex. The results revealed that AL, LT, lens power, AL/CR ratio, and corneal diameter were all significantly correlated with DR (all, P < 0.05). These correlations remained statistically significant when considering VTDR as a dependent variable, with the exception for corneal diameter. However, CC, SE, CCT, and refractive status persistently showed no correlation (all, P > 0.05).

**Table 3.** Associations of ocular biometry and diabetic retinopathy after adjusted for age and sex.

| Age and sex adjusted | Any DR |  | VTDR |  |
| --- | --- | --- | --- | --- |
|  | OR (95%CI) | P-value | OR (95%CI) | P-value |
| Axial length, mm | 0.82 (0.73, 0.91) | **<0.001** | 0.65 (0.52, 0.81) | **<0.001** |
| Corneal curvature, D | 1.05 (0.97, 1.14) | 0.238 | 1.04 (0.91, 1.19) | 0.571 |
| ACD, mm | 0.73 (0.53, 1.00) | 0.052 | 0.49 (0.26, 0.92) | 0.027 |
| Len thickness, mm | 1.52 (1.06, 2.18) | **0.024** | 2.42 (1.27, 4.63) | **0.007** |
| Lens power, D | 0.99 (0.99, 1.00) | **0.003** | 0.99 (0.98, 1.00) | **0.001** |
| AL/CR ratio | 0.24 (0.10, 0.61) | **0.003** | 0.03 (0.00, 0.20) | **<0.001** |
| Spherical equivalent, D | 1.04 (0.99, 1.09) | 0.160 | 1.09 (0.99, 1.21) | 0.080 |
| Corneal diameter, mm | 0.75 (0.57, 0.99) | **0.041** | 0.64 (0.39, 1.03) | 0.068 |
| CCT, μm | 1.00 (0.997, 1.00) | 0.760 | 1.00 (0.996, 1.01) | 0.388 |
| Refractive status |  |  |  |  |
| Emmetropia | Ref |  | Ref |  |
| Hyperopia | 1.01 (0.69, 1.47) | 0.976 | 0.89 (0.46, 1.72) | 0.736 |
| Mild myopia | 0.80 (0.45, 1.44) | 0.456 | 0.46 (0.15, 1.44) | 0.182 |
| Moderate myopia | 0.86 (0.42, 1.76) | 0.676 | 0.49 (0.11, 2.21) | 0.351 |
| High myopia | 1.07 (0.76, 1.51) | 0.710 | 1.20 (0.66, 2.19) | 0.542 |

DR=diabetic retinopathy; VTDR=vision threatened DR; OR=odds ratio; 95%CI=95% confidential interval; ACD=anterior chamber depth; CCT=central corneal thickness; AL/CR ratio=axial length to corneal curvature ratio.
Bold indicates statistical significance.

Table 4 shows the results of further adjusting for other potential confounding factors. Any DR was independently associated with AL (OR = 0.84 per 1 mm increase, 95%CI: 0.74, 0.94), lens power (OR = 0.9951 per 1 D increase, 95%CI: 0.9904, 0.9998), and AL/CR ratio (OR = 0.26 per 1 increase, 95%CI: 0.10-0.70). The ACD only showed a negative correlation with DR when taking it as a quantile. The CC, corneal diameter, and SE were not significantly correlated with presence of DR. Similarly, the presence of VTDR was significantly related to AL (OR = 0.67 per 1 mm increase, 95%CI: 0.54-0.85), lens power (OR = 0.99 per 1 D increase, 95%CI: 0.98, 0.997), and AL/CR ratio (OR = 0.04 per 1 increase, 95%CI: 0.01, 0.25) after adjusting for potential confounding factors. The ACD only showed a negative correlation with VTDR when taking it as a quantile. The CC, corneal diameter, and refractive status were not significantly correlated with presence of VTDR. Figure 1 shows the results of restrictive cubic spline regression analysis evaluating the association between AL and DR. As expect, the odds ratio for any DR and VTDR all tended to decrease as the AL lengthened.

**Table 4.** Multivariable adjusted models to evaluate the associations of ocular biometry and diabetic retinopathy.

| Multivariable<br>adjusted* | Any DR |  | VTDR |  |
| --- | --- | --- | --- | --- |
|  | OR (95%CI) | P-value | OR (95%CI) | P-value |
| **AL** |  |  |  |  |
| Quantile 1 | 1.00 (Ref) |  | 1.00 (Ref) |  |
| Quantile 2 | 0.84 (0.60, 1.18) | 0.324 | 0.83 (0.47, 1.45) | 0.513 |
| Quantile 3 | 0.54 (0.38, 0.78) | **0.001** | 0.73 (0.41, 1.30) | 0.280 |
| Quantile 4 | 0.49 (0.33, 0.70) | **<0.001** | 0.35 (0.18, 0.68) | **0.002** |
| Per 1-mm<br>increase | 0.84 (0.74, 0.94) | **0.003** | 0.67 (0.54, 0.85) | **0.001** |
| **Corneal curvature** |  |  |  |  |
| Quantile 1 | 1.00 (Ref) |  | 1.00 (Ref) |  |
| Quantile 2 | 1.21 (0.85, 1.71) | 0.293 | 0.91 (0.51, 1.61) | 0.744 |
| Quantile 3 | 1.32 (0.93, 1.88) | 0.126 | 0.98 (0.55, 1.76) | 0.943 |
| Quantile 4 | 1.17 (0.81, 1.69) | 0.393 | 0.96 (0.53, 1.74) | 0.883 |
| Per 1-D increase | 1.03 (0.95, 1.12) | 0.466 | 1.02 (0.89, 1.17) | 0.783 |
| **ACD** |  |  |  |  |
| Quantile 1 | 1.00 (Ref) |  | 1.00 (Ref) |  |
| Quantile 2 | 0.58 (0.41, 0.82) | **0.002** | 0.62 (0.35, 1.12) | 0.112 |
| Quantile 3 | 0.65 (0.46, 0.92) | **0.016** | 0.74 (0.42, 1.31) | 0.302 |
| Quantile 4 | 0.58 (0.40, 0.83) | **0.003** | 0.36 (0.18, 0.71) | **0.003** |
| Per 1-mm<br>increase | 0.79 (0.56, 1.10) | 0.158 | 0.56 (0.29, 1.07) | 0.080 |
| **Lens thickness** |  |  |  |  |
| Quantile 1 | 1.00 (Ref) |  |  |  |
| Quantile 2 | 1.14 (0.79, 1.64) | 0.483 | 2.03 (1.10, 3.75) | 0.023 |
| Quantile 3 | 0.92 (0.62, 1.35) | 0.661 | 1.37 (0.68, 2.74) | 0.378 |
| Quantile 4 | 1.31 (0.90, 1.92) | 0.161 | 1.81 (0.90, 3.62) | 0.096 |
| Per 1-mm<br>increase | 1.32 (0.89, 1.95) | 0.168 | 2.03 (1.04, 3.97) | 0.039 |
| **Lens power** |  |  |  |  |
| Quantile 1 | 1.00 (Ref) |  |  |  |
| Quantile 2 | 0.87 (0.61, 1.23) | 0.418 | 0.71 (0.41, 1.24) | 0.230 |
| Quantile 3 | 0.78 (0.55, 1.11) | 0.174 | 0.53 (0.29, 0.97) | **0.039** |
| Quantile 4 | 0.66 (0.45, 0.96) | **0.031** | 0.50 (0.26, 0.95) | **0.033** |
| Per 1-D increase | 0.9951(0.9904,<br>0.9998) | **0.042** | 0.99 (0.98, 0.997) | **0.007** |
| **AL/CR ratio** |  |  |  |  |
| Quantile 1 | 1.00 (Ref) |  |  |  |
| Quantile 2 | 0.84 (0.60, 1.18) | 0.309 | 0.61 (0.35, 1.06) | 0.079 |
| Quantile 3 | 0.62 (0.44, 0.89) | **0.009** | 0.59 (0.34, 1.04) | **0.070** |
| Quantile 4 | 0.65 (0.46, 0.92) | **0.017** | 0.35 (0.19, 0.66) | **0.001** |
| Per 1 increase | 0.26 (0.10, 0.70) | **0.007** | 0.04 (0.01, 0.25) | **0.001** |
| **Refractive status** |  |  |  |  |
| Emmetropia | 1.00 (Ref) |  |  |  |
| Hyperopia | 1.13 (0.75, 1.71) | 0.556 | 1.04 (0.52, 2.07) | 0.913 |
| Mild myopia | 0.75 (0.40, 1.42) | 0.379 | 0.48 (0.15, 1.54) | 0.218 |
| Moderate myopia | 0.87 (0.40, 1.89) | 0.730 | 0.48 (0.10, 2.27) | 0.358 |
| High myopia | 1.14 (0.78, 1.66) | 0.510 | 1.27 (0.67, 2.39) | 0.462 |
| Per 1-D increase | 1.04 (0.99, 1.10) | 0.133 | 1.10 (0.99, 1.23) | 0.064 |
| **Corneal diameter** |  |  |  |  |
| Quantile 1 | 1.00 (Ref) |  |  |  |
| Quantile 2 | 1.27 (0.90, 1.80) | 0.175 | 0.79 (0.44, 1.42) | 0.430 |
| Quantile 3 | 1.00 (0.70, 1.44) | 0.991 | 0.92 (0.52, 1.64) | 0.778 |
| Quantile 4 | 0.92 (0.64, 1.33) | 0.670 | 0.68 (0.37, 1.24) | 0.211 |
| Per 1-mm increase | 0.82 (0.60, 1.11) | 0.205 | 0.72 (0.44, 1.20) | 0.207 |

\*Adjusted for age, sex, duration of diabetes mellitus, HbA1c, total cholesterol, serum creatinine, serum uric acid, body mass index (BMI), and mean blood pressure. DR=diabetic retinopathy; VTDR=vision threatened DR; OR=odds ratio; 95%CI=95% confidential interval; ACD=anterior chamber depth; CCT=central corneal thickness; AL/CR ratio=axial length to corneal curvature ratio. Bold indicates statistical significance.

**Figure 1.**
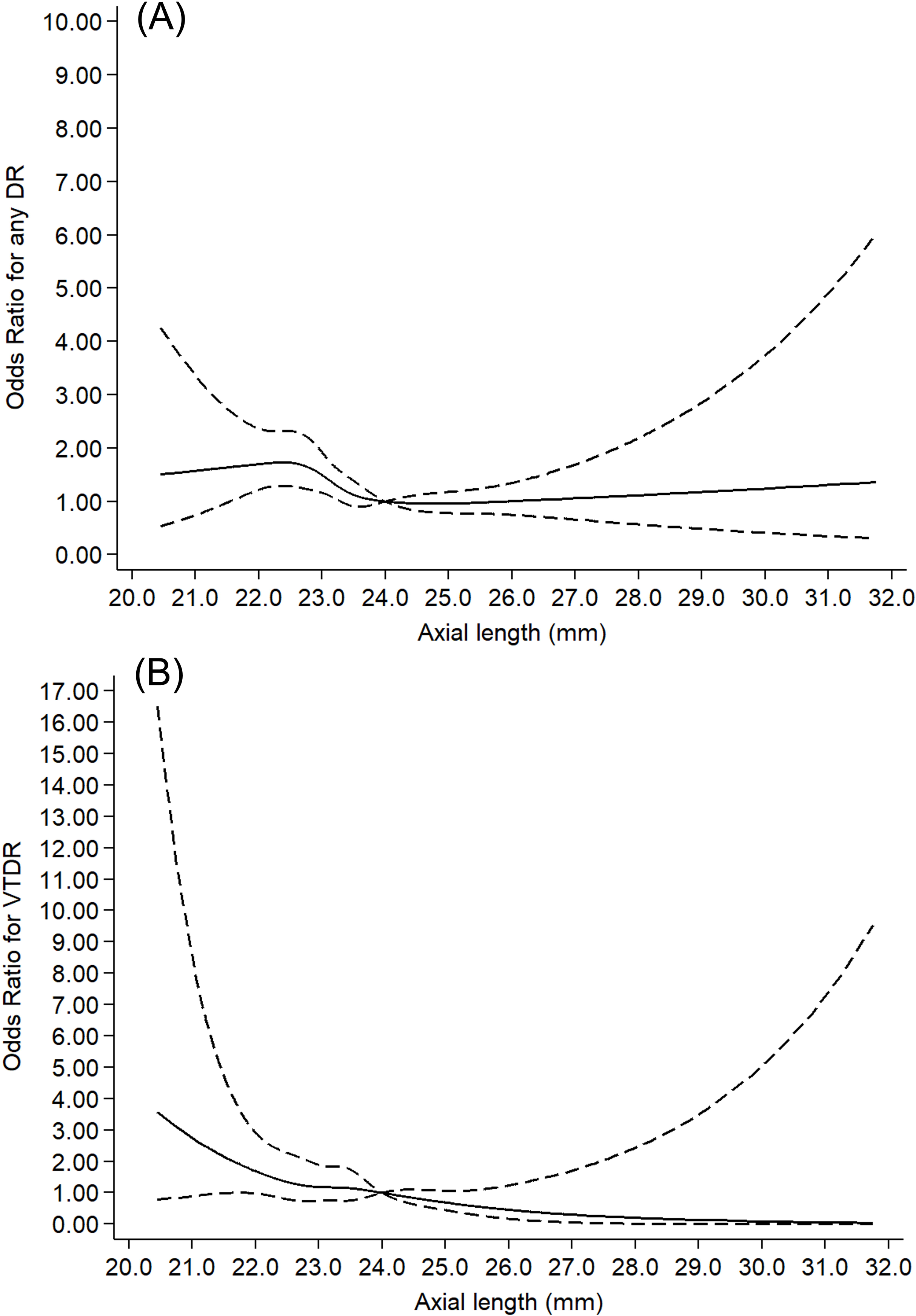
Graph showing the odds ratios for any diabetic retinopathy (DR) and vision threatened DR (VTDR) according to axial length. Data were fitted using a Logistic regression model adjusting for age, sex, duration of diabetes mellitus, HbA1c, total cholesterol, serum creatinine, serum uric acid, BMI, and mean blood pressure. AL was modeled using restricted cubic splines (solid line is the point estimate and dashed lines are 95% confidence limits) with 5 knots at 5%, 27.5%, 50%, 72.55, and 95% percentiles of AL distribution. The reference value is 24.0 mm.

## Discussion

The DR and myopia have been increasing in prevalence in recent decades, and both contribute greatly to visual impairment. Diabetes has been linked to changes in refractive errors under hyperglycaemic conditions. It was reported that high myopia may decrease the progression of DR, even though it is associated with serious ocular complications, such as an increased risk of glaucoma, cataract, and retinal detachment.^17^ Myopia was highly related to the changes of ocular structure, however, which component of the ocular biometry play the major role in this relationship remains unclear. This study demonstrated that longer AL, deeper ACD, higher lens power, and higher AL/CR ratio may all be protective factors against DR and VTDR, independent of age, sex and other potentially confounding factors. However, the CC, corneal diameter, LT, refractive status or SE were not associated with DR. To the best of our knowledge, this is the first study to investigate the ocular components and DR risk in ocular treatment naïve patients with T2DM in the Chinese population.

This study found that refractive status was not associated with presence of DR. Although several small sample clinical and epidemiological studies have suggested that myopia could be a protective factor against DR, this conclusion remains controversial. Moss et al. (1994) conducted a cohort study in 1210 young diabetic patients but reported no correlation of myopia with DR and PDR in univariate analysis, while it was found that myopia may delay the progression from DR to PDR after controlling for confounding factors.^18^ Furthermore, Dogru et al. (1998) reported that high myopia may be a protective factor against PDR in a small sample size retrospective study.^19^ Bazzazi et al. (2017) compared two eyes in anisometropia and verified that high myopia could decrease the incidence of DR, and higher myopia and longer AL provided a greater protective effect.^20^ Several studies based on the Chinese, Korean, and Singaporean population suggest that myopia is protective against PDR, but how different myopia status could influence DR was not mentioned in these studies.^10, 21, 22^ A recent longitudinal cohort study demonstrated that refractive status did not influence the incidence and progression of DR. Consistent with this cohort study, the present study indicate that different myopia status may not influence DR risk.

Longer AL was associated with lower risk for both DR and VTDR, which was consistent with previous studies. Several population-based studies suggested that AL played a different role in DR genesis and development in different ethnicities, although some contradictory results existed in literature.^7–9^ A recent cohort study of Singaporean population demonstrated that the any DR risk decreased by 42% for each 1 mm increase. However, the aforementioned study reported no correlation of AL with the risk of VTDR.^6^ Considering the high prevalence of myopia in the juvenile population, it is worth keeping a watchful eye on how the incidence of DR in this generation changes over time. Further longitudinal studies with large sample size are needed.

The mechanism of the protective effect of longer AL against DR remains unclear. Several factors may play a role in this protective phenomenon. First, it was might related to the pathological alteration caused by AL that increases with the progression of myopia. This may result in a thinner retina and choroid as well as reduced blood flow in the retina.^23, 24^ The low perfusion status relatively decreases the structural damage of the retinal vessel wall, and also prevents biochemical damage caused by high glycogen accumulation. Second, oxygen demand is also decreased as the retina becomes thinner, which alleviates the retina’s hypoxic status in diabetic patients.^25^ Third, posterior vitreous detachment (PVD) and synchesis may occur as myopia progresses, which enables the retina to gain oxygen from the liquefied vitreous body, resulting in a decreased rate of angiogenesis.^26^ Fourth, alterations in cytokines could also be a potential mechanism, such as vascular endothelial growth factor, pigment epithelium-derived factor, tumor necrosis factor, erythropoietin, and TGF-β.^27^ Further basic studies are warranted to elaborate the underling mechanism.

Both AL and corneal radius are closely related to the refractive status, with the finding that AL/CR is linearly dependent on the diopter in populations aged 40 to 64. It was also reported that AL/CR had a stronger relationship with myopia compared to other ocular biometry parameters such as AL, ACD and CC. Previous only 1 study have evaluated the influence of AL/CR ratio and lens power on risk of DR and reported that both AL/CR and lens power were related to DR, which is consistent with our results. These findings indicated that lens power and corneal refractive components also play a role in protective effects of ocular elongation against DR.

Few studies have investigated the relationship between DR and other biometrical parameters including CC, ACD, and LT. Pierro et al.^28^ found that the LT increased in patients with insulin-dependent diabetes and the thicker LT was associated with lower risk for PDR. Another hospital-based study did not observed any correlation between LT and DR after adjusting confounding factors.^29^ The population-based Beijing Eye Study reported that ACD was not related to presence of DR.^11^ We found that the deeper ACD was associated with lower risk of DR in when taking it as a quantile. Thus, further studies are required to verify our finding that a correlation may exist between ACD and lens thickness and DR or VTDR.

The strengths of our study include the enrolment of only ocular treatment naïve patients with T2DM, a relatively large sample size based on the community population, and fully adjusting for confounding factors. This study also has several limitations. First, the causal relationship between biometry parameters and DR could not be determined due to the inherent features of a cross-sectional study, which need to be verified in a longitudinal study. Second, the subjects in this study were all type 2 diabetic patients, and the conclusion of this study needs to be confirmed in further studies with type 1 diabetic patients. Finally, the subjects were all recruited from communities in south China. Considering myopia has an ethnic heterogeneity, the generalisation of the conclusions is limited. Multi-ethnic and multi-centre studies are warranted to verify our findings.

## Conclusions

This study demonstrated that longer AL, deeper ACD, higher lens power, and higher AL/CR ratio may be protective factors against DR, independent of age, sex and other potentially confounding factors. Further studies are warranted to elaborate the potential mechanisms of how the ocular biometry alterations influence DR. Considering the high prevalence of myopia in the juvenile population, it may prove beneficial to pay attention to how the incidence of DR in this generation changes over time.

## Acknowledgments

This study was supported by the National Natural Science Foundation of China (81570843; 81530028; 81721003). The funding organizations had no role in the design or conduct of the study; collection, management, analysis, and interpretation of the data; preparation, review, or approval of the manuscript; and decision to submit the manuscript for publication.

## Author Contributions

WW and WH had full access to all the data in the study and take responsibility for the integrity of the data and the accuracy of the data analysis. Study concept and design: WW, WL, MH, WH. Acquisition, analysis, or interpretation of data: LS, YL, XG, XK, WL. Drafting of the manuscript: LS, WW. Critical revision of the manuscript for important intellectual content: All authors. Statistical analysis: WW. Obtained funding: WH. Administrative, technical, or material support: MH, WW. Study supervision: MH.

